# Chronic fentanyl induces sex-specific anxiolytic-like effects in opioid-dependent rats

**DOI:** 10.64898/2026.10.19.760254

**Authors:** Lara Caglayan, Fiona Zeyl, Enzo Gamba, Marcelo Febo, Adriaan Bruijnzeel

## Abstract

Fentanyl is used as a treatment for severe pain, and it is a major contributor to opioid misuse and overdose fatalities. While it has been established that opioids are widely misused for their euphoric effects, little is known about the effects of chronic fentanyl exposure on anxiety-like behavior. We determined the effects of chronic fentanyl on anxiety-like behavior, locomotor activity, dependence, and body weight in adult male and female Wistar rats (n = 8 per sex and treatment) that received saline or fentanyl (0.6 mg/kg/day) via minipumps. Locomotor activity was measured in a small open field (SOF) before pump implantation. After implantation, rats were retested in the SOF, evaluated for anxiety-like behavior in the elevated plus-maze (EPM) and large open field (LOF), and it was determined if they were opioid dependent by counting somatic withdrawal signs. Fentanyl increased locomotor activity in the SOF and the LOF. In the EPM, fentanyl increased open-arm time in males but not in females. In the LOF, fentanyl increased entries into, and shortened the latency to enter, the anxiogenic inner zones, most prominently in females. Naloxone precipitated a somatic withdrawal syndrome in fentanyl-treated rats of both sexes, and fentanyl prevented body weight gain in males and attenuated it in females. To provide insight into the behavior across the LOF and EPM, an additional correlational analysis was conducted. In saline-treated males, but not females, anxiety-related measures were correlated between the two tests. Rats that made more open-arm entries in the EPM made more entries into, and spent more time in, the center zone of the LOF (r = 0.78 and 0.75), and the percentage of time on the open arms correlated with time spent in the middle zone of the LOF (r = 0.73). In conclusion, fentanyl decreases anxiety-like behavior in opioid dependent animals. These findings have significant implications as they suggest that dependent individuals might continue to misuse fentanyl for its anxiolytic effects. Furthermore, patients treated with chronic fentanyl for severe pain and comorbid anxiety might continue to benefit from the drug’s anxiolytic effects.

## 1. Introduction

Opioid use disorder (OUD) remains a critical public health crisis in the United States, driven increasingly by illicit synthetic opioids rather than the prescription medications that historically fueled the epidemic (Ciccarone, 2021). Recent national data indicate that approximately 8.9 million individuals aged 12 or older misused opioids in 2023, and 5.7 million met the criteria for an opioid use disorder (SAMHSA, 2024). Concurrently, the overdose crisis has been fundamentally altered by the proliferation of illicitly manufactured fentanyl. Total drug overdose deaths escalated dramatically over the last decade, peaking at approximately 112,000 in the 12-month period ending August 2023, before declining to approximately 67,000 in the 12-month period ending April 2026 (Ahmad et al., 2026). Synthetic opioids other than methadone, predominantly fentanyl, continue to account for more than half of all overdose deaths (Ahmad et al., 2026).

Fentanyl is a highly lipophilic, full agonist at the mu-opioid receptor that is approximately 100 times more potent than morphine, has a faster onset of action, and produces tolerance and physical dependence with repeated exposure (Comer and Cahill, 2019). Fentanyl is also widely used clinically, as an anesthetic adjunct and for the treatment of severe and chronic pain, and in the latter setting it is often administered continuously, for example by transdermal patches that release the drug over several days, so that exposure is sustained rather than fluctuating with each dose (Comer and Cahill, 2019). Because fentanyl has largely replaced heroin and prescription opioids in the illicit drug supply (Mattson et al., 2021; DEA, 2024), preclinical studies that characterize the behavioral effects of fentanyl directly, rather than extrapolating from morphine or heroin, are needed to understand the factors that maintain fentanyl misuse.

Anxiety disorders are among the most prevalent psychiatric conditions and are closely associated with opioid use. In humans, endogenous opioid signaling reduces anxiety; this is supported by the observation that opioid receptor antagonists increase anxiety and panic symptoms (Colasanti et al., 2011). Consistent with this, acute administration of the full mu-opioid agonist hydromorphone in humans attenuates the cortisol response to acute psychosocial stress (Bershad et al., 2018). Anxiety and depressive disorders are common among people who use prescription opioids (Sullivan et al., 2006; Feingold et al., 2017), and longitudinal data indicate that pre-existing mood and anxiety disorders increase the risk of nonmedical prescription opioid use and opioid use disorder (Martins et al., 2009; Martins et al., 2012).

Consistent with these observations, mu-opioid receptor agonists reduce anxiety-like behavior in rodents. Acute morphine increases open-arm exploration in the elevated plus-maze (EPM) (Koks et al., 1999; Rezayof et al., 2009), acute oxycodone decreases anxiety-like behavior in the EPM in rats of both sexes, with a larger effect in males (Bruijnzeel et al., 2022), and chronic once-daily fentanyl injections decreased anxiety-like behavior in the open field and the EPM in mice (Fujii et al., 2019). However, the direction of these effects depends heavily on the dose, the pattern of exposure, and the state of the animal (Roeska et al., 2008; Ahmadi et al., 2023). Whether chronic fentanyl decreases anxiety-like behavior in opioid-dependent rats, whether such an effect generalizes across tests, and whether it differs between males and females have not been established.

The relationship between opioids and anxiety is complicated by the fact that the presence and the absence of the drug produce opposing affective states. In rats, continuous fentanyl administration via osmotic minipumps induces a dependent state in which spontaneous or naloxone-precipitated withdrawal is accompanied by somatic signs, elevations in brain reward thresholds, and loss of body weight (Bruijnzeel et al., 2006; Bruijnzeel et al., 2007; Liu et al., 2008). Opiate withdrawal also produces anxiogenic-like effects in the EPM (Schulteis et al., 1998), and current models of opioid addiction posit that the negative emotional state of withdrawal becomes a major driver of continued drug use (Koob, 2020). The net effect of fentanyl on anxiety-like behavior is therefore expected to depend on whether animals are tested under steady-state drug conditions or during withdrawal. This underscores the need to characterize anxiety-like behavior in animals whose level of dependence has been investigated in the same study.

Opioids also alter locomotor activity. Fentanyl and its analogs stimulate locomotor activity in rodents (Varshneya et al., 2019; Gaulden et al., 2021). Because the measures of anxiety-like behavior obtained in exploration-based tests are sensitive to overall activity (Rodgers and Dalvi, 1997), locomotor activity has to be assessed alongside anxiety-like behavior in order to interpret drug effects. Furthermore, sex differences have been reported in the analgesic, reinforcing, and motoric effects of opioids (Craft, 2008; Becker and Chartoff, 2019), in fentanyl self-administration and relapse-like behavior (Townsend et al., 2019; Bakhti-Suroosh et al., 2021), and in the anxiolytic-like effect of oxycodone (Bruijnzeel et al., 2022), so that it is critical to include males and females in the studies.

The EPM and the large open field (LOF) are among the most widely used and best validated tests of anxiety-like behavior in rodents. In both tests, avoidance of the exposed regions of the apparatus, the open arms of the EPM and the center of the LOF, is interpreted as anxiety-like behavior (Pellow et al., 1985; Prut and Belzung, 2003; Walf and Frye, 2007), and clinically effective anxiolytics increase exploration of these regions in maze-naive animals (Pellow et al., 1985; Pellow and File, 1986; File, 1993). The two tests capture overlapping but not identical behavioral constructs; we previously showed in a pooled group of adult male and female Wistar rats that behavioral parameters are correlated both within and between the EPM and the LOF and that both tests are sensitive to sex (Knight et al., 2021). Examining both tests in the same animals, together with the spatial distribution of their behavior, therefore makes it possible to determine whether a drug effect generalizes across tests of anxiety-like behavior or is specific to one apparatus.

The goal of the present study was to determine the effects of chronic fentanyl on anxiety-like behavior, locomotor activity, and dependence in adult male and female rats. Rats received saline or fentanyl (0.6 mg/kg/day) via osmotic minipumps. In contrast to intermittent injections, which because of the short duration of action of fentanyl produce a daily cycle of intoxication and withdrawal, minipumps provide continuous delivery that resembles the sustained exposure produced by transdermal fentanyl in patients and induces a stable state of dependence (Bruijnzeel et al., 2006; Liu et al., 2008). Locomotor activity was assessed in a small open field (SOF) before and after pump implantation, anxiety-like behavior was assessed in the EPM and the LOF, physical dependence was assessed by measuring naloxone-precipitated somatic withdrawal signs, and body weight was recorded throughout the post-implantation period. We hypothesized that chronic fentanyl would decrease anxiety-like behavior in both tests and that fentanyl would increase locomotor activity and decrease body weight gain.

## 2. Materials and methods

### 2.1. Animals

Wistar rats (males 200-225 g, females 175-200 g, Charles River, Raleigh, NC) were socially housed in a climate-controlled vivarium on a reversed 12 h light-dark cycle (light off at 7 AM). Food and water were available ad libitum. All animal procedures were performed in accordance with the University of Florida Institutional Animal Care and Use Committee as well as the National Institutes of Health guidelines.

### 2.2. Experimental design

Adult male and female Wistar rats (n = 32; 16 males and 16 females) were studied under an identical schedule, yielding four groups defined by pump treatment (saline or fentanyl) and sex (n = 8 per group). After arrival, all rats were handled and weighed for at least one week. Baseline locomotor activity was then assessed in three SOF sessions (15 min each) conducted before pump implantation; the groups that were subsequently assigned to saline or fentanyl were compared on these baseline measures and did not differ (see Results). On day 0, rats were implanted subcutaneously with an osmotic minipump that delivered either saline or fentanyl (0.6 mg/kg/day). The fentanyl dose was based on prior studies in which we investigated the development of opioid dependence (Bruijnzeel et al., 2007; Liu et al., 2008). Behavioral testing resumed after implantation of the pumps. Testing was conducted according to the following schedule: a post-pump SOF session on day 3–4, the EPM on day 4–5, assessment of naloxone-precipitated somatic withdrawal signs on two consecutive days (day 5–7), and the LOF on day 13–14. Body weight was recorded throughout the post-surgical period. On each somatic-sign test day, rats received saline or naloxone (0.03 mg/kg, s.c.) 10 min before a 10-min scoring period. The naloxone dose was based on our prior opioid dependence studies (Bruijnzeel et al., 2006; Bruijnzeel et al., 2007; Liu et al., 2008). Between the somatic-sign tests and the LOF test, rats underwent one functional brain imaging session under anesthesia on day 9–11. The imaging data are beyond the scope of the present report and will be described in a separate paper.

### 2.3. Drugs

Fentanyl citrate and naloxone hydrochloride were purchased from Sigma-Aldrich (St. Louis, MO, USA) and dissolved in sterile saline (0.9% sodium chloride). Fentanyl and naloxone doses are expressed as the salt. Carprofen (Levafen 50 mg/ml; Pivetal) was purchased from Patterson Veterinary Supply (Ocala, FL, USA).

### 2.4. Osmotic minipump implantation

To induce dependence, rats were prepared with osmotic minipumps (Alzet model 2ML2, 14-day pumps; Durect Corporation, Cupertino, CA) that delivered fentanyl. Fentanyl was administered subcutaneously (s.c.) for 14 days at a dose of 0.6 mg/kg/day (fentanyl citrate salt; 0.38 mg/kg/day fentanyl base). The concentration of the pump solution was adjusted to the body weight of each rat on the day of implantation. Control rats were prepared with pumps that delivered vehicle (saline). The pumps were filled under aseptic conditions with sterile syringes and solutions and were implanted s.c. in an aseptic manner under isoflurane/oxygen anesthesia (1–3% isoflurane). Carprofen (5 mg/kg, s.c.) was administered once, immediately before pump implantation, to provide perioperative analgesia. All rats also received naloxone (0.03 mg/kg, s.c.) immediately before pump implantation as a precaution against respiratory depression during the combination of isoflurane anesthesia and the onset of fentanyl delivery. The flank was shaved and disinfected with 70% alcohol and betadine, a 1.5-cm incision was made posterior to the rib cage, a subcutaneous pocket large enough to accommodate the pump was created with sterile blunt scissors, the pump was inserted, and the incision was closed with non-absorbable sutures or staples. The sutures and staples were removed approximately 10 days after surgery.

### 2.5. Small open field test

The SOF test was conducted to assess locomotor activity in a dark room with the lights turned off (Febo et al., 2003; Bruijnzeel et al., 2016). The total distance traveled was measured with an automated animal activity cage system (AccuScan Instruments, Columbus, OH, USA). The system consisted of four animal activity cages made of clear acrylic (40 cm × 40 cm × 30 cm; L x W x H), with 16 equally spaced (2.5 cm) infrared beams across the length and width of the cage. The infrared beams were positioned 2 cm above the cage floor and were connected to a VersaMax analyzer which sent information to a computer that displayed beam data through VersaDat software. The test setup was cleaned with a Nolvasan solution between animals.

### 2.6. Elevated plus-maze test

The elevated plus-maze test is used to assess anxiety-like behavior and was conducted as described previously (Rylkova et al., 2009; Qi et al., 2016; Bruijnzeel et al., 2019). The test apparatus consisted of four black polypropylene arms (Coulbourn Instruments, Whitehall, PA). The two "open" arms had 0.5 cm ledges, and the two "closed" arms had 30 cm walls. The open arms were placed opposite of each other. The arms were 10 cm wide, 50 cm long, and were placed on 55 cm tall acrylic legs. Testing occurred in a quiet, dimly lit (75 lux) room. At the beginning of each test, the rats were placed on an open arm facing the center of the apparatus. The rats were allowed to explore the apparatus for 5 min, and their behavior was recorded with a camera mounted above the maze. In some trials, video tracking was started before the rat had been placed on the maze and was stopped too early. Therefore, the portion of each recording preceding placement was discarded, and each recording was trimmed at the end so that the analyzed period ran from the moment the rat was placed on the maze to 260 s thereafter and was the same for every animal in the study. The EPM was divided into five zones (two open arms, two closed arms, and a center zone). Behavior was analyzed automatically (center-point detection) using EthoVision XT 13 software (Noldus Information Technology, Leesburg, VA). The following behavioral parameters were automatically determined: duration and entries into the open arms, closed arms, and center zone, and total distance traveled. The percentage of open-arm entries (open-arm entries/total arm entries × 100) and the percentage of time on the open arms (time on open arms/total time on the arms × 100) were calculated. Occupancy heatmaps were generated from the raw center-point coordinates (Section 2.9). The apparatus was cleaned with a Nolvasan solution between rats.

### 2.7. Somatic signs of naloxone-precipitated withdrawal

Somatic signs of opioid withdrawal were assessed on two consecutive days (days 5–7 after pump implantation). Before the test days, the rats were habituated to the Plexiglas observation chambers (26 × 26 × 26 cm; L × W × H) for 5 min per day on three consecutive days. On each test day, rats received an injection of saline or naloxone (0.03 mg/kg, s.c.), with the order of the two injections counterbalanced across rats, and 10 min later they were placed individually in the observation chamber, in which somatic withdrawal signs were recorded for 10 min by an experienced observer who was blind to the treatment conditions (Bruijnzeel et al., 2006; Bruijnzeel et al., 2007; Liu et al., 2008). The following signs were recorded: ptosis, facial fasciculations, abdominal constrictions, body shakes, escape attempts, erection or ejaculation, abnormal posture, diarrhea, and paw tremor. Each sign was scored as the number of occurrences during the 10-min observation period, and a total somatic sign score was calculated for each rat as the sum of all signs. The observation chamber was cleaned with a Nolvasan solution between rats.

### 2.8. Large open field test

The LOF test is used to assess locomotor activity and anxiety-like behavior (Liebsch et al., 1998). The test was conducted for 5 min in a dimly lit room (75 lux), as described previously (Bruijnzeel et al., 2016; Qi et al., 2016; Tan et al., 2019). The LOF apparatus consisted of a large arena measuring 120 x 120 x 60 cm (L x W x H). The arena was made of black high-density polyethylene panels that were fastened together and placed on a plastic bottom plate (Faulkner Plastics, Miami, FL). The rats’ behavior was recorded with a camera mounted above the arena and analyzed with EthoVision XT 13 software (Noldus Information Technology, Leesburg, VA). The LOF was divided into three zones: an outside zone (20 cm wide), a middle zone (20 cm wide), and a center zone (40 x 40 cm; L x W). The following behaviors were analyzed: total distance traveled, distance traveled in each zone (outside, middle, and center), number of entries into each zone, and latency to enter the middle and center zone. Occupancy heatmaps were generated from the raw center-point coordinates (Section 2.9). The LOF was cleaned with a Nolvasan solution between rats.

### 2.9. Occupancy heatmaps

Spatial occupancy heatmaps for the SOF, EPM, and LOF were generated from the raw position data of each animal rather than with the heatmap generators built into the acquisition software. For the SOF, the position of the rat at each sample was derived from the AccuScan beam-break coordinate export (VersaDat .DXY files), which records the first and last interrupted infrared beams in the X and Y directions; the position was taken as the center of this bounding box, and the 2.5-cm beam spacing was used to convert beam indices to centimeters. For the elevated plus-maze and large open field, the X and Y coordinates of the center point of the rat (cm) were exported from EthoVision XT 13 as raw track data (25–30 samples/s). The full post-pump 15-min SOF, the 260-s EPM record, and the 5-min LOF record were used. For each animal, the time elapsed between successive samples was accumulated into square spatial bins (1.25 cm for the SOF; 2 cm for the EPM and LOF), lightly smoothed with a Gaussian kernel (σ = 1 bin), and expressed as the percentage of the session time spent in each bin. The LOF coordinates were scaled per cohort to the 120 × 120 cm arena. Group maps were calculated as the mean of the eight individual maps of each sex × drug group and are displayed on a common color scale within each test (square-root scaling; upper limit set at the 99.5th percentile of the four group maps). All processing was done in Python 3 using NumPy, pandas, and openpyxl for data handling, SciPy (ndimage) for smoothing, and Matplotlib for plotting.

### 2.10. Statistical analysis

In the SOF test, total distance traveled was analyzed both over the full 15-min session and for the first 5-min block alone. Pre-pump activity across the three baseline sessions (days 1–3) was analyzed with a three-way mixed-model analysis of variance (ANOVA) with sex (male, female) and drug (saline, fentanyl) as between-subjects factors and session (day 1, day 2, day 3) as a within-subjects factor. Sphericity was assessed with Mauchly’s test; because it was satisfied, uncorrected degrees of freedom are reported (Greenhouse–Geisser correction was applied where sphericity was violated). Post-pump implantation activity was compared between groups with a two-way (sex × drug) analysis of covariance (ANCOVA), using each animal’s mean day 1–3 activity as the covariate. Homogeneity of regression slopes, homogeneity of variance (Levene’s test), and normality of residuals (Shapiro–Wilk test) were confirmed. Adjusted (estimated marginal) means were computed at the grand-mean covariate value.

Somatic signs were counted and because the signs were not normally distributed and were zero-inflated, they were analyzed with nonparametric tests. For the total somatic sign score and for each of the nine individual signs, differences among the four groups (saline pump/saline injection, saline pump/naloxone injection, fentanyl pump/saline injection, and fentanyl pump/naloxone injection) were assessed with Kruskal–Wallis tests, followed by Dunn’s post-hoc comparisons with Bonferroni correction. Sex differences were evaluated with Mann–Whitney U tests. Because no sex differences were detected, males and females were combined for the analyses reported.

Elevated plus-maze measures were analyzed with two-way ANOVAs with drug (saline, fentanyl) and sex (male, female) as between-subjects factors. Significant interactions were followed by Bonferroni-corrected post-hoc comparisons. Large open field measures were analyzed with two-way ANOVAs with drug (saline, fentanyl) and sex (male, female) as between-subjects factors. Significant interactions were followed by Bonferroni-corrected post-hoc comparisons.

In the control (saline-treated) animals, relationships between behavioral measures across tests were examined with Pearson correlations among the EPM, LOF, and SOF parameters, computed separately for males and females (n = 8 per sex) and displayed as correlation heatmaps. These correlational analyses were exploratory and were not corrected for multiple comparisons.

Body weight was expressed as a percentage of each animal’s weight on the first post-surgical day (day 1 = 100%) to correct for the mass of the implanted osmotic pump. Post-surgical trajectories (days 5–13) were analyzed with a mixed-model ANOVA with drug (saline, fentanyl) and sex as between-subjects factors and day as a within-subjects factor, and separately within each sex (drug × day). Greenhouse–Geisser correction was applied where Mauchly’s test indicated a violation of sphericity, and significant interactions were followed by Bonferroni-corrected comparisons of the two drug groups at each day.

All statistical tests were two-tailed, and effects were considered significant at α = 0.05. Group sizes were n = 8 per sex × drug group (n = 16 per sex) for all analyses unless stated otherwise. Analyses of variance and covariance were performed in IBM SPSS Statistics Version 32 (IBM Corp., Armonk, NY, USA). Correlations and bar graphs were prepared in GraphPad Prism Version 11 (GraphPad Software, San Diego, CA, USA). Occupancy heatmaps, correlation matrices, and the processing of the raw tracking data were carried out in Python Version 3.11 using the NumPy (Harris et al., 2020), SciPy (Virtanen et al., 2020), pandas (McKinney, 2010) and Matplotlib (Hunter, 2007) libraries.

## 3. Results

### 3.1. Small open field

#### 3.1.1. Baseline (days 1–3)

During the pre-pump baseline, full-session total distance declined across the three sessions (session, F(2,56) = 7.58, p = 0.001), and this decline was greater in females than males (session × sex interaction, F(2,56) = 4.44, p = 0.016); females were also more active overall (main effect of sex, F(1,28) = 6.17, p = 0.019). Females traveled farther than males on day 1 (p = 0.006) and day 2 (p = 0.020) but not on day 3 (Bonferroni-corrected comparisons), and in females activity on day 3 was lower than on day 1 (p = 0.011). Importantly, there was no main effect of drug (F(1,28) = 0.18, p = 0.673) and no interaction of drug with session or sex, confirming that the groups did not differ in locomotor activity before pump implantation (Figure 1; Table S1). When the analysis was restricted to the first 5-min block of each baseline session, no effect reached significance.

**Figure 1.**
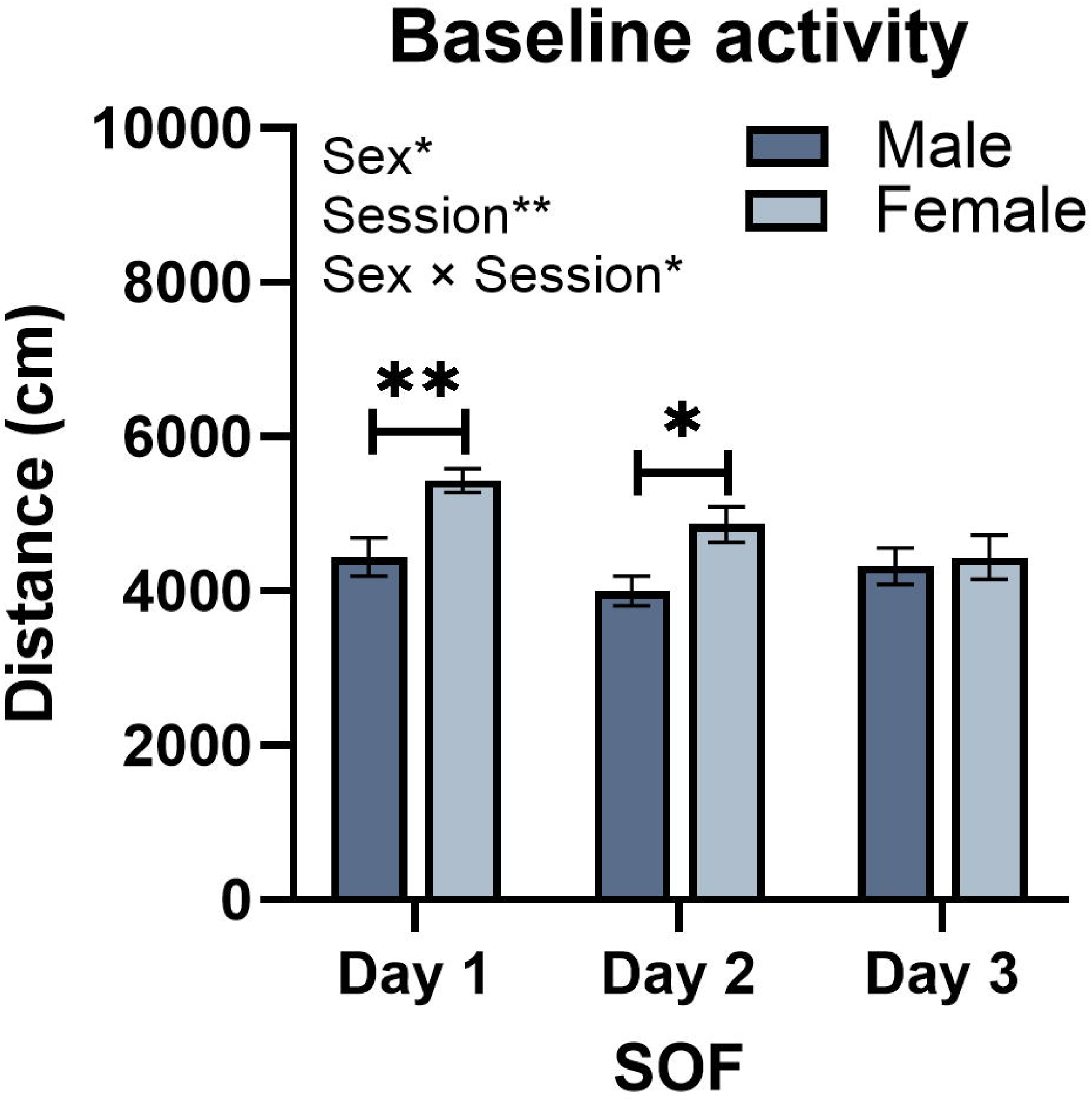
Baseline locomotor activity in the small open field before implanting the minipumps. Total distance traveled (cm) during the three 15-min baseline sessions (days 1–3) in male and female rats (bars: mean ± SEM; n = 16 per sex; saline and fentanyl groups are pooled because treatment had not yet started and the groups did not differ). Data were analyzed by mixed-model ANOVA (sex × session); significant main effects and the interaction are indicated in the top-left corner (Sex, Session, Sex × Session; *p < 0.05, **p < 0.01). Activity declined across sessions (session, p = 0.001), more steeply in females than in males (session × sex, p = 0.016). Females traveled farther than males on day 1 (**p < 0.01) and day 2 (*p < 0.05; Bonferroni-corrected comparisons within day, indicated by brackets) but not on day 3.

#### 3.1.2. Post-pump full 15-minute session

After adjustment for baseline, post-pump total distance did not differ significantly between drug groups over the full session (Figure 2A; Table S2; F(1,27) = 3.69, p = 0.065). Baseline activity significantly predicted post-pump activity, F(1,27) = 5.44, p = 0.027. Neither the main effect of sex nor the sex × drug interaction was significant.

**Figure 2.**
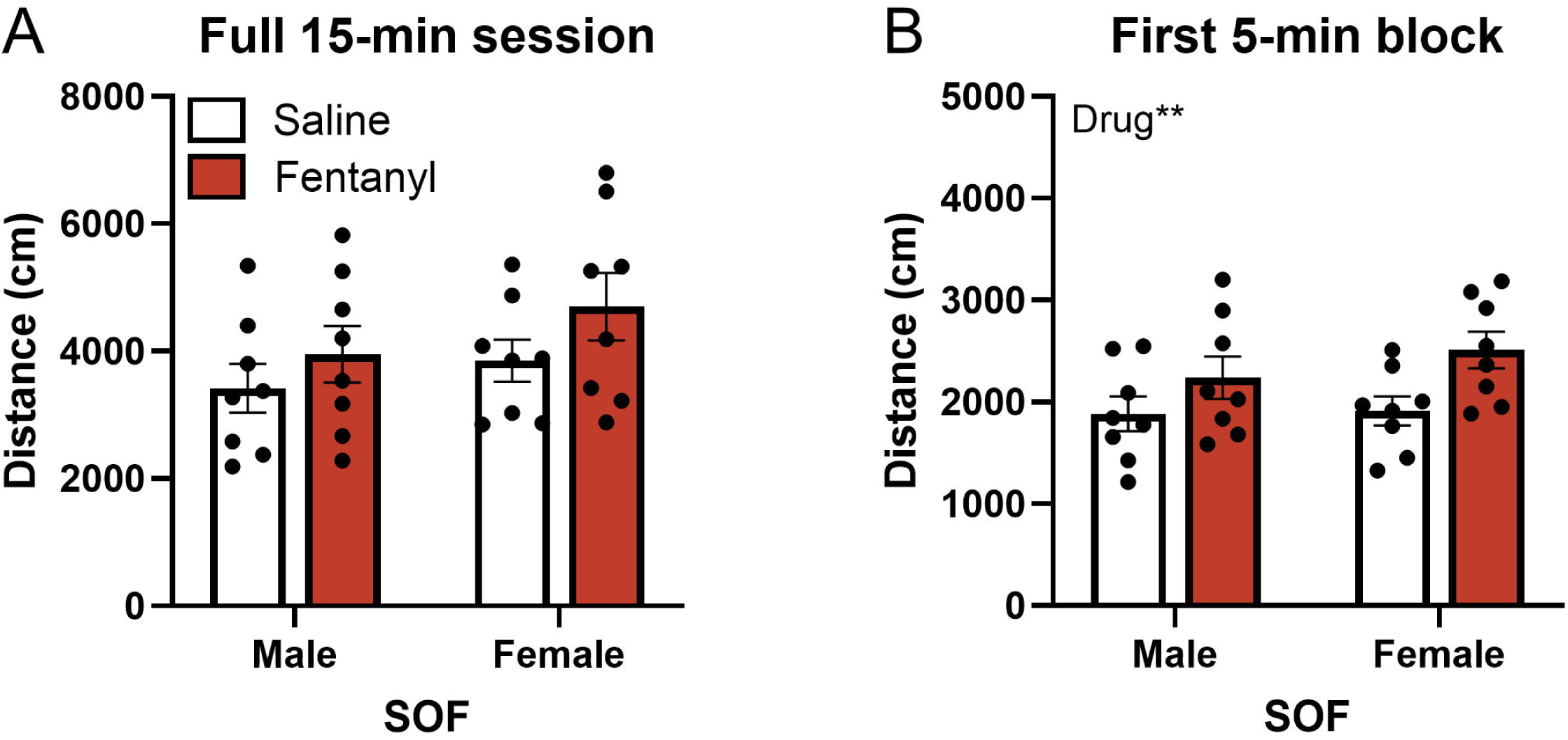
Locomotor activity in the small open field test after minipump implantation. Total distance traveled during the post-pump session over (A) the full 15-min session and (B) the first 5-min block, by sex and drug treatment (bars: mean ± SEM; circles: individual animals; n = 8 per group). Data were analyzed by two-way (sex × drug) ANCOVA with each animal’s mean pre-pump activity (days 1–3) as the covariate. The main effect of drug was significant for the first 5-min block (**p < 0.01, fentanyl vs saline; indicated in the top-left corner of panel B) but not for the full session (p = 0.065); neither the main effect of sex nor the sex × drug interaction was significant.

#### 3.1.3. Post-pump first 5-minute block

When activity was restricted to the first 5-min block (Figure 2B; Table S2), a significant main effect of drug emerged: fentanyl-treated animals traveled farther than controls, F(1,27) = 8.28, p = 0.008. The baseline covariate was not predictive in this window (F(1,27) = 1.21, p = 0.281), and the drug effect was comparable in an unadjusted ANOVA (p = 0.012). The main effect of sex and the sex × drug interaction were not significant.

### 3.2. Somatic signs of naloxone-precipitated withdrawal

Naloxone precipitated somatic withdrawal signs in fentanyl-treated rats (Figure 3; Table 1). The total somatic sign score differed among the four groups (Kruskal–Wallis H(3) = 37.4, p < 0.001). Fentanyl-treated rats that received naloxone displayed more somatic signs (26.6 ± 2.0) than fentanyl-treated rats that received saline (1.2 ± 0.3), saline-pump rats that received naloxone (1.1 ± 0.3), or saline-pump rats that received saline (2.1 ± 0.5; Dunn’s post-hoc tests, all p < 0.001). At the level of individual signs, the Kruskal–Wallis test was significant for eight of the nine signs. Dunn’s post-hoc comparisons showed that fentanyl/naloxone rats displayed more ptosis, facial fasciculations, abdominal constrictions, body shakes, escape attempts, abnormal posture, diarrhea, and erection/ejaculation than saline/naloxone rats (all Bonferroni-corrected p < 0.05); paw tremor did not differ among the groups (p = 0.26). Males and females did not differ in the total somatic sign score or in any individual sign (Mann–Whitney U tests, all p > 0.05), and the data were therefore combined across sex.

**Figure 3.**
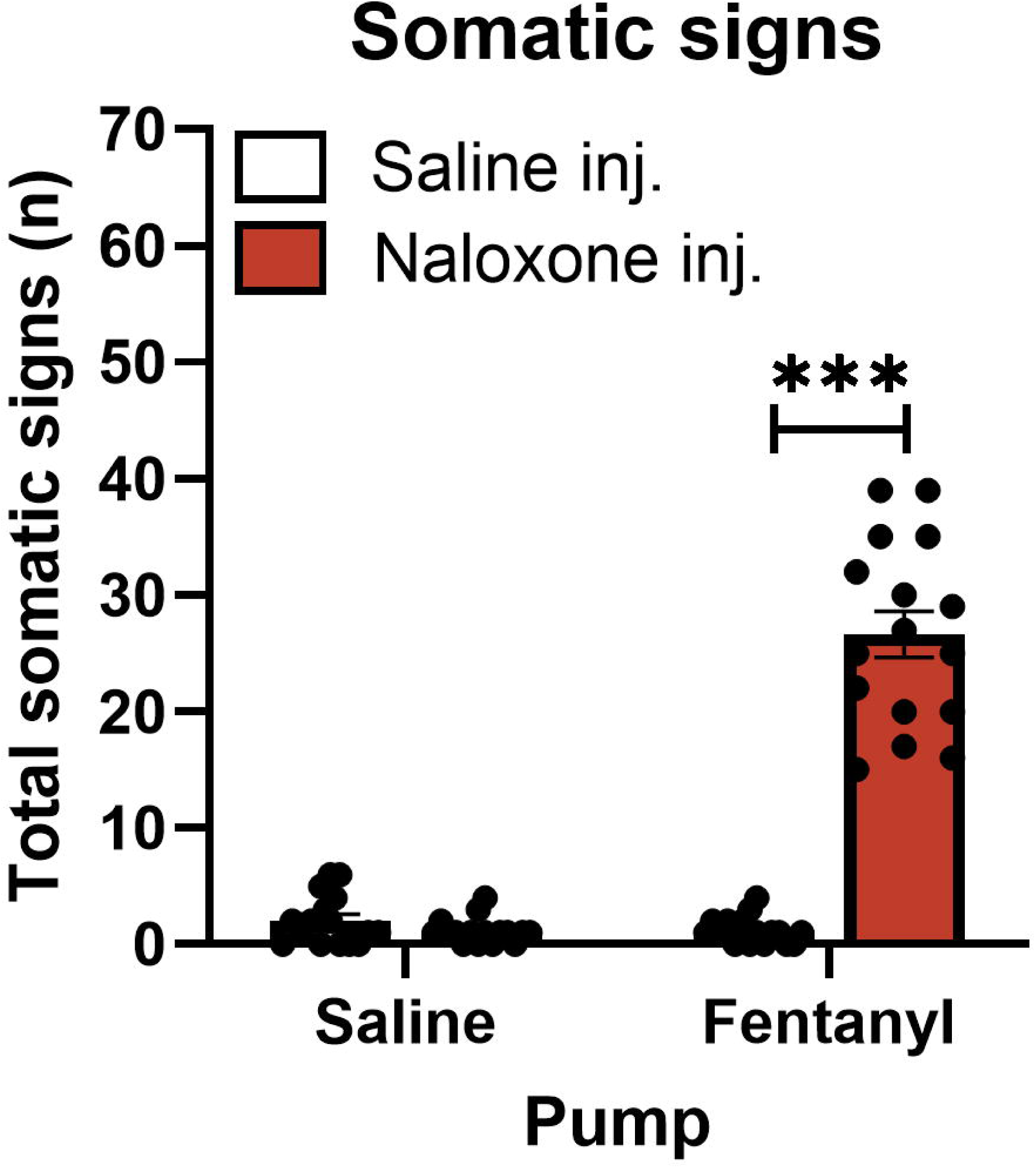
Total somatic signs of naloxone-precipitated withdrawal. Total number of somatic withdrawal signs (mean ± SEM; n = 16 per group, 8 male and 8 female) in saline- and fentanyl-treated rats that received a saline or naloxone injection 10 min before a 10-min observation period. ***p < 0.001 versus all other groups (Kruskal–Wallis with Dunn’s Bonferroni-corrected post-hoc tests). Individual signs are reported in Table 1.

**Table 1.** Somatic signs of naloxone-precipitated withdrawal, by group.

| Somatic sign | Saline pump +<br>Saline | Saline pump +<br>Naloxone | Fentanyl pump<br>+ Saline | Fentanyl pump<br>+ Naloxone |
| --- | --- | --- | --- | --- |
| Ptosis | 0.25 ± 0.17 | 0.44 ± 0.27 | 0.00 ± 0.00 | 3.19 ± 0.57*** |
| Facial | 0.62 ± 0.20 | 0.25 ± 0.11 | 0.56 ± 0.16 | 6.62 ± 0.66*** |
| Abdominal | 0.44 ± 0.20 | 0.06 ± 0.06 | 0.31 ± 0.15 | 2.19 ± 0.54*** |
| Shakes | 0.19 ± 0.10 | 0.25 ± 0.11 | 0.25 ± 0.14 | 5.69 ± 0.77*** |
| Escape | 0.56 ± 0.34 | 0.00 ± 0.00 | 0.06 ± 0.06 | 4.69 ± 1.58*** |
| Erection/ejaculation | 0.00 ± 0.00 | 0.00 ± 0.00 | 0.00 ± 0.00 | 0.44 ± 0.22* |
| Abnormal posture | 0.00 ± 0.00 | 0.12 ± 0.09 | 0.00 ± 0.00 | 2.75 ± 0.60*** |
| Diarrhea | 0.00 ± 0.00 | 0.00 ± 0.00 | 0.00 ± 0.00 | 0.88 ± 0.09*** |
| Paw tremor | 0.00 ± 0.00 | 0.00 ± 0.00 | 0.00 ± 0.00 | 0.19 ± 0.14 |
| **Total somatic signs** | **2.06 ± 0.54** | **1.12 ± 0.27** | **1.19 ± 0.29** | **26.62 ± 1.99\*\*\*** |

Group means ± SEM (n = 16 per group; 8 males and 8 females). Rats received a saline or naloxone injection and somatic signs were scored during the observation period. Asterisks denote a significant difference from the Saline pump + Naloxone group (Kruskal–Wallis with Dunn’s Bonferroni-corrected post-hoc test): \*p < 0.05, \*\*p < 0.01, \*\*\*p < 0.001.

### 3.3. Elevated plus-maze

In the EPM, chronic fentanyl increased open-arm exploration in a sex-dependent manner (Figure 4). There were significant drug × sex interactions for the percentage of time on the open arms (F(1,28) = 7.84, p = 0.009) and for the time spent on the open arms (F(1,28) = 8.41, p = 0.007). Bonferroni-corrected post-hoc comparisons indicated that fentanyl-treated males spent a greater percentage of time on the open arms than saline-treated males (29.5 ± 2.7% vs 11.8 ± 4.2%, p < 0.001) and more time on the open arms (75.3 ± 6.8 s vs 29.9 ± 10.5 s, p < 0.001), whereas fentanyl- and saline-treated females did not differ (p = 0.96 and p = 0.996, respectively).

**Figure 4.**
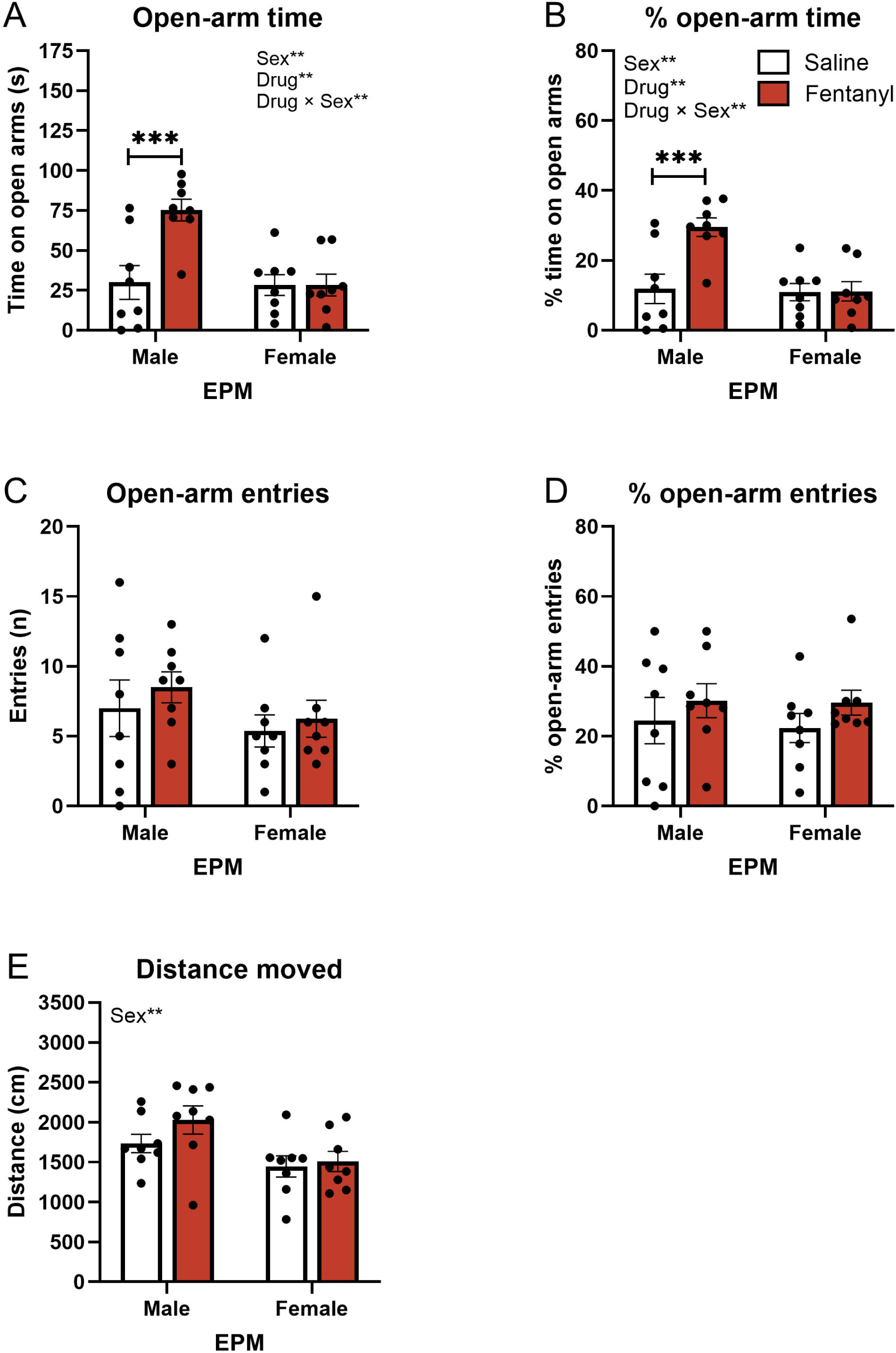
Anxiety-like behavior in the elevated plus-maze. (A) Time on the open arms, (B) percentage of time on the open arms, (C) open-arm entries, (D) percentage of open-arm entries, and (E) distance traveled, by sex and drug treatment (bars: mean ± SEM; circles: individual animals; n = 8 per group). Data are from the 260-s EPM recordings. Data were analyzed by two-way (drug × sex) ANOVA; significant main effects and interactions are indicated in the top-left corner of each panel (Sex, Drug, Drug × Sex; **p < 0.01). Where the drug × sex interaction was significant, Bonferroni-corrected post-hoc comparisons are indicated by brackets (***p < 0.001, fentanyl vs saline in males).

In contrast, the number of open-arm entries and the percentage of open-arm entries showed no drug × sex interaction and no main effect of drug (all p ≥ 0.20). Distance traveled showed a main effect of sex, with males traveling a greater distance than females (F(1,28) = 8.34, p = 0.007), but no main effect of drug (F(1,28) = 1.61, p = 0.215) and no interaction, indicating that the increased open-arm exploration of fentanyl-treated males was not secondary to a general change in locomotor activity.

### 3.4. Large open field

In the LOF, chronic fentanyl increased locomotor activity and increased exploration of the inner zones (Figure 5). Fentanyl-treated rats traveled a greater total distance than saline-treated rats (main effect of drug, F(1,28) = 19.87, p < 0.001), with no effect of sex (F(1,28) = 1.84, p = 0.186) and no interaction (F(1,28) = 0.03, p = 0.871). Fentanyl also increased entries into the outer zone (drug, F(1,28) = 23.98, p < 0.001; females made more entries than males, sex F(1,28) = 30.80, p < 0.001) and into the middle zone (drug, F(1,28) = 15.15, p < 0.001), with a drug × sex interaction (F(1,28) = 4.41, p = 0.045). Bonferroni-corrected comparisons showed that fentanyl increased middle-zone entries in females (18.9 ± 1.7 vs 6.8 ± 0.8, p < 0.001) but not in males (p = 0.22).

**Figure 5.**
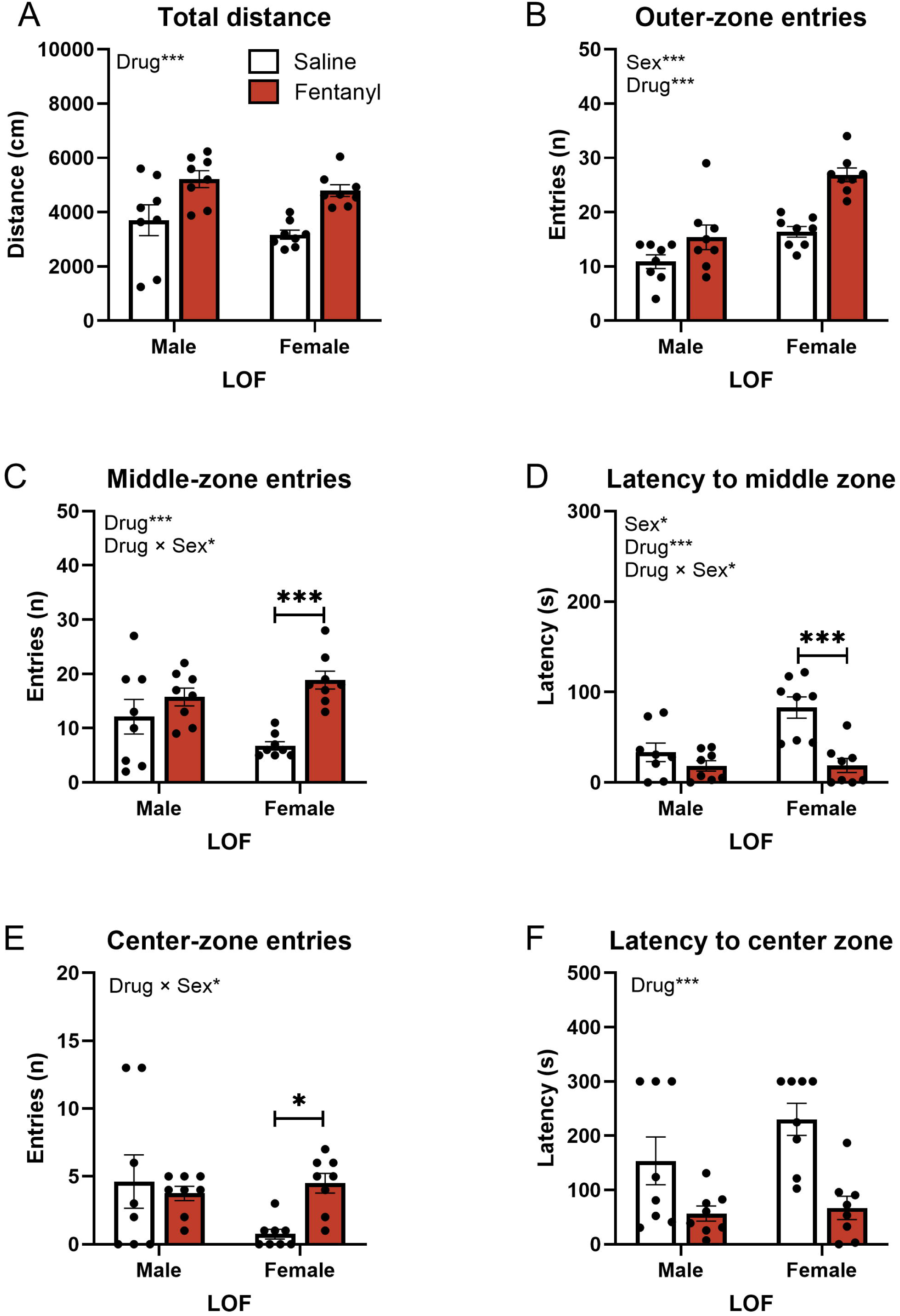
Behavior in the large open field. (A) Total distance traveled, (B) entries into the outer zone, (C) entries into the middle zone, (D) latency to enter the middle zone, (E) entries into the center zone, and (F) latency to enter the center zone, by sex and drug treatment (bars: mean ± SEM; circles: individual animals; n = 8 per group). Rats that never entered a zone were assigned the maximum latency (300 s). Data were analyzed by two-way (drug × sex) ANOVA; significant main effects and interactions are indicated in the top-left corner of each panel (Sex, Drug, Drug × Sex; *p < 0.05, ***p < 0.001). Where the drug × sex interaction was significant, Bonferroni-corrected fentanyl-versus-saline comparisons within each sex are indicated by brackets (*p < 0.05, ***p < 0.001).

Fentanyl also shortened the latency to enter the inner zones. The latency to enter the middle zone was reduced by fentanyl (drug, F(1,28) = 18.54, p < 0.001), and a drug × sex interaction (F(1,28) = 7.10, p = 0.013; sex, F(1,28) = 7.40, p = 0.011) indicated that this reduction was most pronounced in females (Bonferroni-corrected fentanyl vs saline: 18.9 ± 7.9 s vs 82.9 ± 11.8 s, p < 0.001; males, p = 0.26), which showed the longest latencies under saline. Similarly, fentanyl reduced the latency to enter the center zone (drug, F(1,28) = 19.56, p < 0.001). Center-zone entries showed a drug × sex interaction (F(1,28) = 4.46, p = 0.044), with fentanyl increasing center-zone entries in females (Bonferroni-corrected 4.5 ± 0.7 vs 0.8 ± 0.4, p = 0.022) but not males (p = 0.58) and no significant main effects (both p > 0.15). Together, these results indicate that chronic fentanyl increased locomotor activity and promoted exploration of the inner zones of the LOF, an effect that was generally stronger in females. The spatial distribution of these effects in all three tests is shown in the group-averaged occupancy maps (Figure 6).

**Figure 6.**
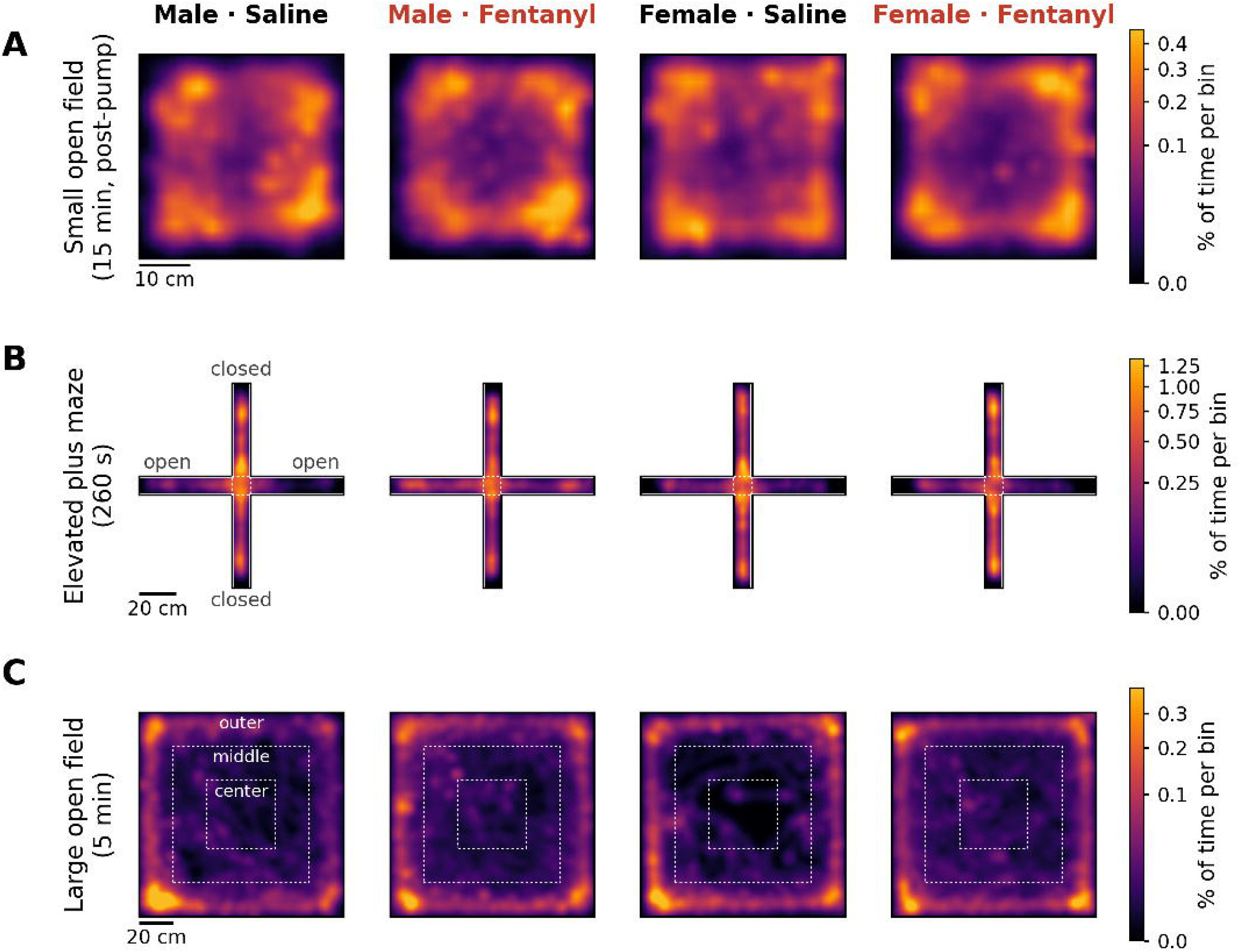
Spatial occupancy heatmaps in the small open field, elevated plus-maze, and large open field. Time-weighted occupancy maps (percentage of session time per spatial bin) averaged across the eight rats of each group (male saline, male fentanyl, female saline, female fentanyl). (A) Small open field, full 15-min post-pump session (40 × 40 cm arena; 1.25-cm bins). (B) Elevated plus-maze, 260-s session (2-cm bins); closed arms are oriented vertically and open arms horizontally, and the dashed square marks the center platform. (C) Large open field, 5-min session (120 × 120 cm arena; 2-cm bins); the dashed squares mark the middle (80 × 80 cm) and center (40 × 40 cm) zones. The color scale is shared within each row (square-root scaling; upper limit at the 99.5th percentile of the row). Scale bars: 10 cm (A) and 20 cm (B, C). n = 8 per group.

### 3.5. Cross-test behavioral correlations

In control (saline-treated) animals, individual differences in behavior were correlated across tests in a sex-dependent manner: these cross-test relationships were present in males but absent in females (Figures 7 and 8). In males, the strongest correlations were between locomotor activity in the first SOF session (15-min total distance) and open-arm exploration in the EPM: this baseline distance predicted the percentage of open-arm entries (r = 0.90), open-arm entries (r = 0.81), and the percentage of time and the time spent on the open arms (both r = 0.77; all p ≤ 0.024). Locomotor measures were also related between the EPM and the LOF: distance traveled in the EPM correlated with distance traveled in the LOF (r = 0.85, p = 0.008), with greater exploration of the inner (middle) zone (middle-zone entries, r = 0.81; middle-zone duration, r = 0.85), less time in the outer zone (r = −0.77), and with LOF center-zone entries (open-arm entries vs. center-zone entries, r = 0.78). By contrast, SOF distance did not correlate with any LOF parameter.

**Figure 7.**
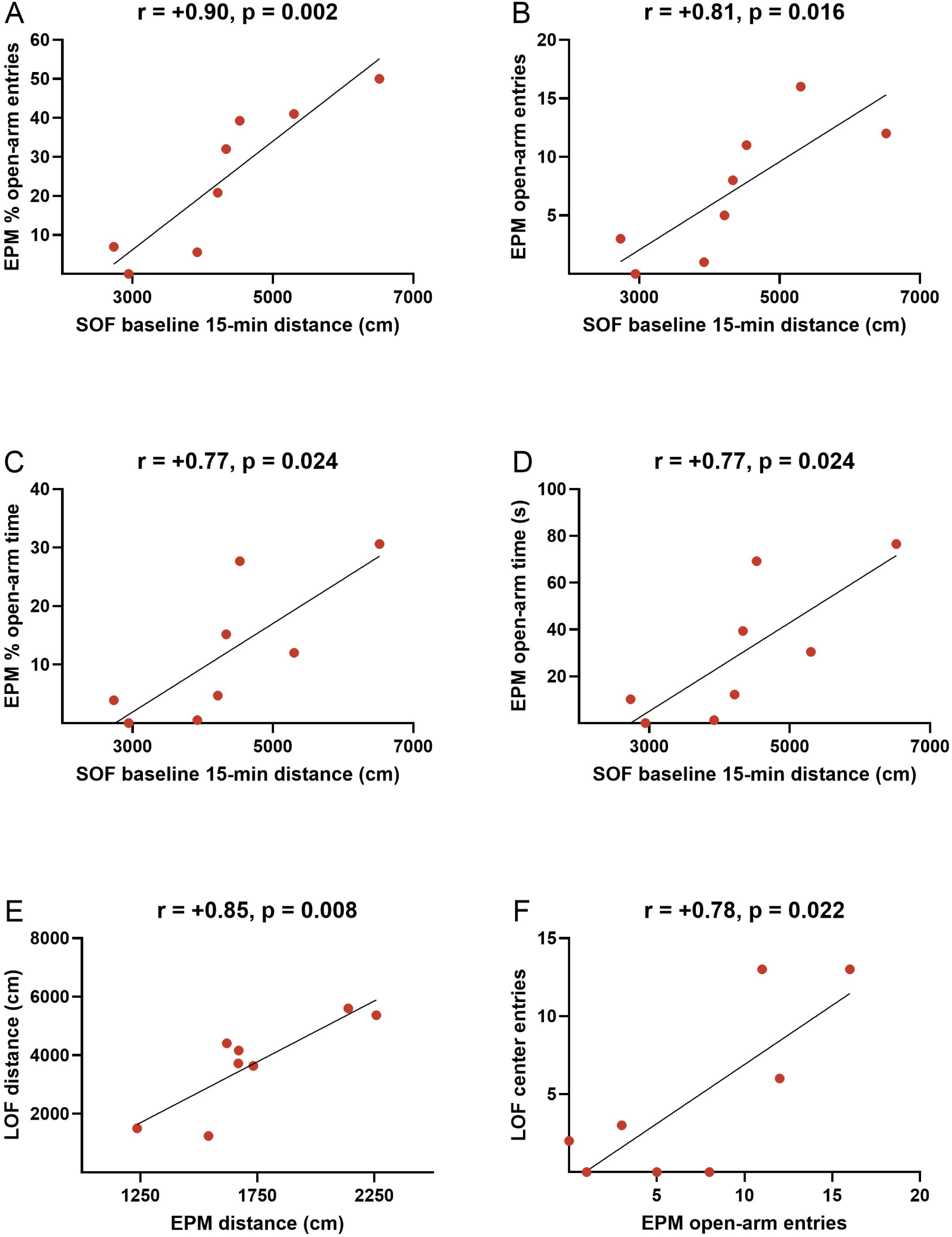
Cross-test correlations in saline-treated males. Scatterplots with Pearson correlation coefficients relating small open field and elevated plus-maze measures (A–D) and elevated plus-maze and large open field measures (E–F) in control (saline-treated) males (n = 8).

**Figure 8.**
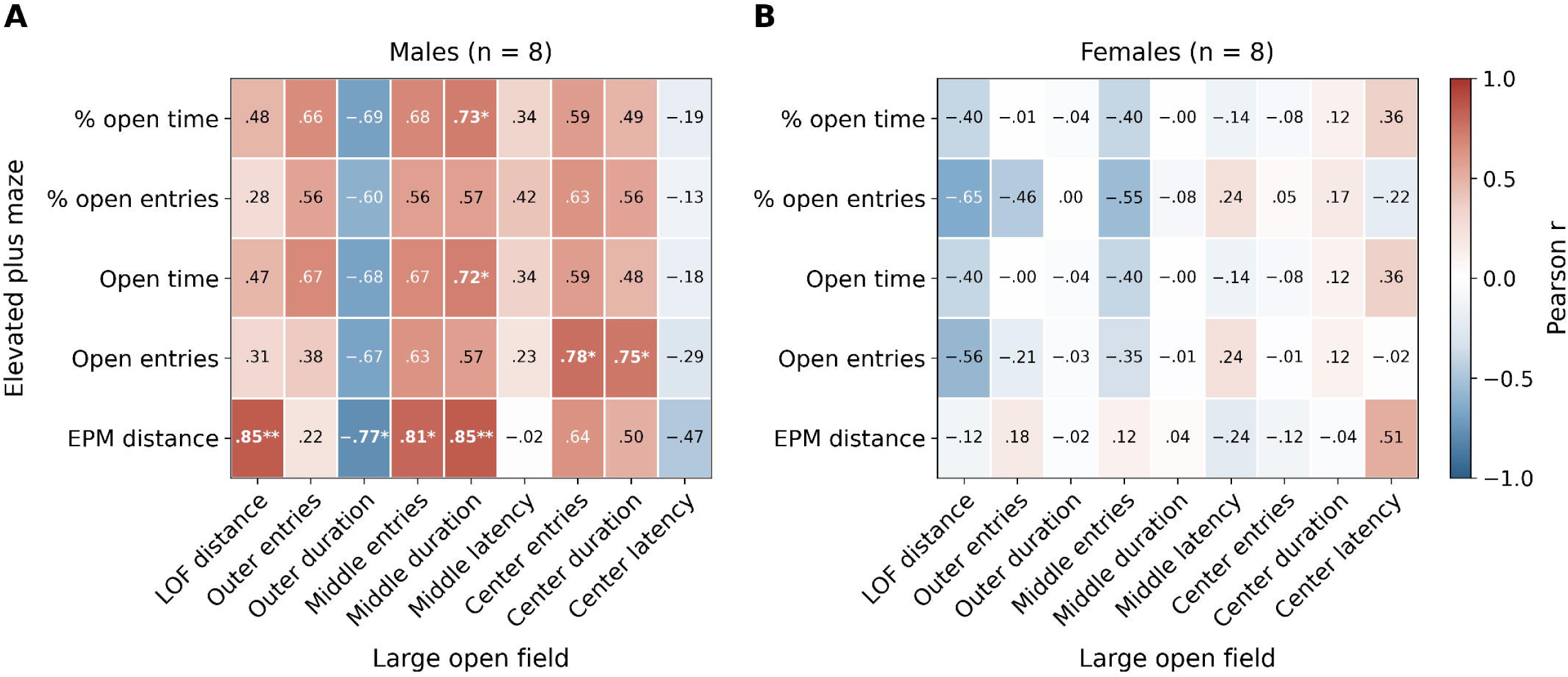
Sex-dependent correlations between elevated plus-maze and large open field behavior. Heatmaps show Pearson correlation coefficients between elevated plus-maze and large open field parameters in control (saline-treated) males (A) and females (B) (n = 8 per sex); cross-test correlations were present in males but essentially absent in females.

In females, none of these cross-test correlations were significant (Figure 8). Thus, individual differences in locomotor and exploratory behavior were consistent across the EPM, LOF, and SOF in males but not in females. Representative cross-test correlations are shown as scatterplots (Figure 7), and the EPM × LOF correlation matrices for males and females are shown as heatmaps (Figure 8).

### 3.6. Body weight

Fentanyl suppressed body-weight gain in a sex-dependent manner (Figure 9). Across both sexes, fentanyl-treated rats weighed less than saline controls (drug main effect F(1,28) = 36.4, p < 0.001) and the two groups diverged progressively over days (drug × day F(7,196) = 6.30, p < 0.001); this divergence itself differed by sex (sex × drug × day F(7,196) = 3.58, p = 0.001). In males, saline rats gained weight steadily (to 108.2 ± 0.8% of day 1 by day 13) whereas fentanyl rats failed to gain weight (97.9 ± 0.9%), yielding large effects of drug (F(1,14) = 79.2, p < 0.001) and a strong drug × day interaction (F(7,98) = 16.5, p < 0.001). Fentanyl males were significantly lighter than saline males from day 5 onward (Bonferroni p < 0.05 at days 5, 6, and 9–13).

**Figure 9.**
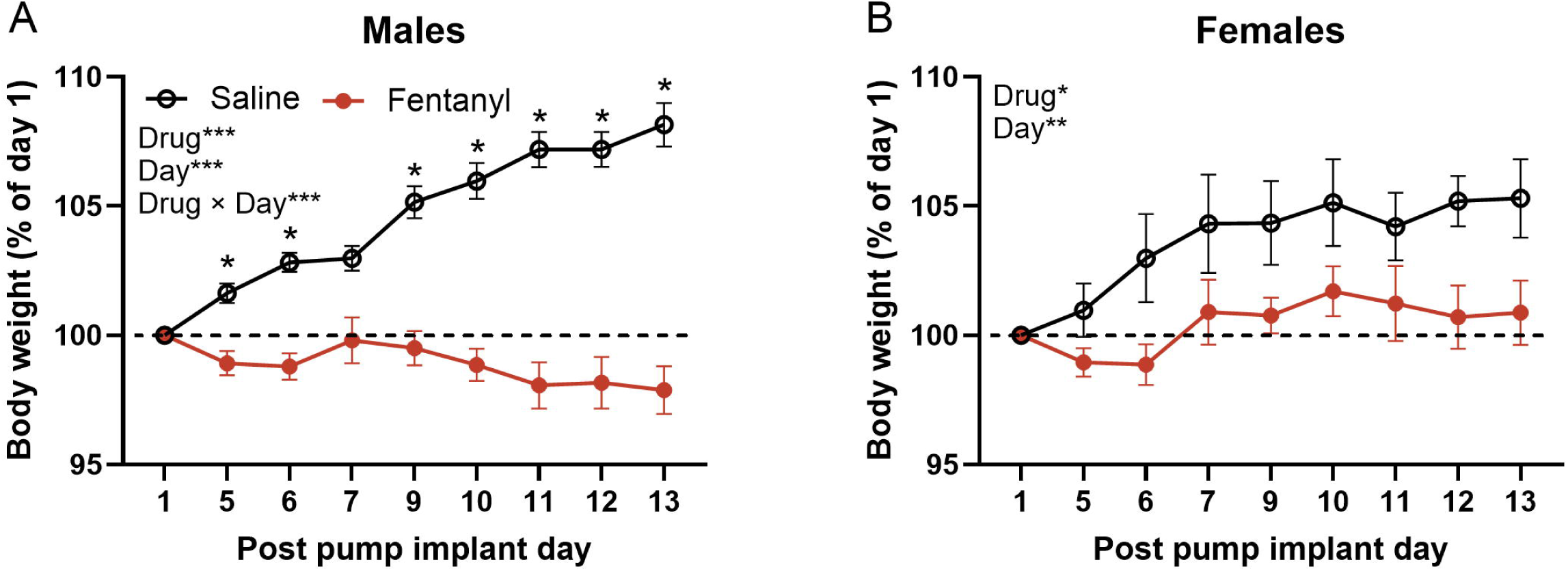
Body weight after minipump implantation. Body weight expressed as a percentage of the first post-surgical day (day 1 = 100%) in males (A) and females (B). Points are group means ± SEM (n = 8 per group) for saline- and fentanyl-treated rats across post-surgery days 1–13. Data were analyzed by mixed-model ANOVA (drug × day) within each sex; significant main effects and interactions are indicated in the top-left corner of each panel (Drug, Day, Drug × Day; *p < 0.05, **p < 0.01, ***p < 0.001); asterisks in (A) mark days on which fentanyl males differed from saline males (Bonferroni-corrected p < 0.05).

In females the drug effect was weaker and did not vary over time: fentanyl females weighed less than saline females (drug F(1,14) = 5.75, p = 0.031) but the two trajectories remained parallel (drug × day F(7,98) = 0.50, p = 0.84, ns). For example, on day 13 the body weight of the saline females was 105.3 ± 1.5% of day 1 weight versus 100.9 ± 1.2% in the fentanyl females.

## 4. Discussion

The present study characterized the effects of chronic fentanyl on anxiety-like behavior, locomotor activity, and physical dependence in adult male and female rats. Chronic fentanyl decreased anxiety-like behavior in both the EPM and the LOF. However, the sex in which the effect was most pronounced differed between the tests. Fentanyl increased open-arm time in the EPM in males, whereas it increased exploration of the inner zones of the LOF most clearly in females. Fentanyl also increased locomotor activity in the SOF and the LOF. Naloxone precipitated a robust somatic withdrawal syndrome in fentanyl-treated rats of both sexes, confirming that the animals were physically dependent at the time of testing. Finally, exposure to fentanyl prevented body weight gain in males and attenuated it in females. Together, these findings indicate that chronic fentanyl, at a dose that produces opioid dependence, has anxiolytic-like and locomotor-stimulant effects that are modulated by sex.

### Fentanyl and anxiety-like behavior in the elevated plus-maze

Chronic fentanyl increased the percentage of time and the absolute time spent on the open arms of the EPM, and this effect was only found in the males. Fentanyl-treated males spent approximately 30% of their time on the open arms, compared with about 10% for saline-treated males, whereas the two female groups did not differ. The anxiolytic-like effect is consistent with the effects of other mu-opioid receptor agonists in this test, including acute morphine (Koks et al., 1999; Rezayof et al., 2009) and acute oxycodone (Bruijnzeel et al., 2022), and with the report that chronic fentanyl decreases anxiety-like behavior in mice (Fujii et al., 2019). Importantly, the effect in males was driven by the time spent on the open arms rather than by the number of open-arm entries or the total distance traveled, neither of which was affected by fentanyl in the EPM. An increase in open-arm time without a change in entries or distance cannot be attributed to hyperactivity and instead indicates that fentanyl-treated males spent more time on the open arms once they entered them. A comparable dissociation, an increase in the percentage of time on the open arms without a change in the percentage of open-arm entries, has been observed with chronic diazepam in the original validation of the test (Pellow et al., 1985). The persistence of an anxiolytic-like effect after several days of continuous fentanyl exposure contrasts with the anxiogenic-like effect that develops over 8 days of twice-daily morphine injections (Ahmadi et al., 2023). While intermittent injections produce significant peaks and troughs in systemic opioid concentrations, continuous minipump delivery maintains stable, steady-state drug exposure.

The male-specific anxiolytic-like effect in the EPM mirrors the sex difference that we previously reported for acute oxycodone, which also decreased anxiety-like behavior more in males than in females (Bruijnzeel et al., 2022). Several factors may contribute to this pattern. Female rats generally display more open-arm exploration and higher activity in the EPM than males (Scholl et al., 2019; Knight et al., 2021), which may leave less room for a further drug-induced reduction in anxiety-like behavior, although in the present study the saline-treated females and males spent a similar percentage of time on the open arms. Sex differences in the pharmacokinetics and pharmacodynamics of opioids, including in receptor density and in gonadal hormone modulation of mu-opioid receptor signaling, are well documented (Craft, 2008; Becker and Chartoff, 2019). Furthermore, the estrous cycle, which was not monitored in the present study, can affect behavior in the EPM (Walf and Frye, 2007). Although these effects are not always observed (Scholl et al., 2019), this unmonitored variable could have contributed to the variability of the female data. A more general implication is that the anxiolytic-like effects of opioids in females may be smaller, or may require different conditions to be detected, than in males.

### Fentanyl and anxiety-like behavior in the large open field

In the LOF, fentanyl-treated rats entered the middle and center zones more often and with shorter latencies than saline-treated rats. These fentanyl effects were largest in females, which under saline conditions were the most avoidant group, with the longest latencies to enter the middle zone and the fewest center entries. Reduced avoidance of the exposed center of an open field is a classical indicator of an anxiolytic-like effect (Prut and Belzung, 2003), and the convergence of reduced avoidance of the open arms of the EPM and of the inner zones of the LOF strengthens the interpretation that chronic fentanyl decreases anxiety-like behavior rather than producing an effect that is confined to a single apparatus.

This interpretation must nevertheless be tempered by the concurrent increase in locomotor activity. Fentanyl increased the total distance traveled in the LOF by approximately 40–50% in both sexes, and measures such as the number of zone entries and the time spent in the center scale with how much an animal moves (Rodgers and Dalvi, 1997; Prut and Belzung, 2003). It is therefore likely that part of the increase in inner-zone entries reflects the increase in activity. Two observations argue that hyperactivity does not account for the entire effect. First, the latency to first enter the middle zone, which depends on the willingness of the animal to leave the wall rather than on the amount of movement thereafter, was strongly reduced by fentanyl. Second, the reduction in anxiety-like behavior in the EPM was not accompanied by an increase in distance traveled. We have therefore treated the LOF parameters as activity-sensitive indices and did not statistically covary for locomotion, as adjusting for a treatment-altered variable would inappropriately remove part of the fentanyl effect under study.

### Relationship between the two tests and the sex-dependent pattern of effects

A notable outcome of the present study is that the anxiolytic-like effect of fentanyl was most evident in males in the EPM and in females in the LOF. Several explanations can be considered. The two tests were conducted at different stages of fentanyl exposure: the EPM on day 4–5 after pump implantation and the LOF on day 13–14. Preclinical literature demonstrates that male and female rodents often develop tolerance to the behavioral and physiological effects of mu-opioid agonists at distinct rates (Craft, 2008; Becker and Chartoff, 2019), raising the possibility that the anxiolytic response in males had already undergone tolerance by the second week of continuous exposure. Alternatively, this divergence may reflect the distinct dimensions of threat captured by the two assays and the fact that these tests do not measure the same constructs in the two sexes: factor analyses indicate that behavior in the EPM is governed primarily by anxiety in male rats but by activity in female rats (Fernandes et al., 1999), and sex differences in anxiety-like behavior depend on the test employed (Johnston and File, 1991; Scholl et al., 2019; Knight et al., 2021). Furthermore, the saline-treated females were the most avoidant group in the LOF but not in the EPM. As a result, the baseline range available to detect a drug-induced reduction in avoidance differed between tests and sexes. Finally, females may be more sensitive to the locomotor-stimulant effects of opioids (Craft, 2008; Gaulden et al., 2021), a factor that would affect the activity-sensitive LOF measures more heavily than the EPM measures.

The cross-test correlations in the saline-treated animals are consistent with this interpretation. In males, the distance traveled in the EPM correlated with the distance traveled and with the exploration of the inner zones in the LOF, and activity in the first SOF session predicted open-arm exploration in the EPM, indicating that individual differences in activity and exploration were consistent across tests. In females, these relationships were absent. This pattern extends our earlier observation that the EPM and the LOF capture overlapping but not identical constructs (Knight et al., 2021) and suggests that in females the two tests may tap partly different traits, in which case test-specific drug effects are not unexpected. Taken together, the findings emphasize that conclusions about sex differences in the anxiolytic-like effects of opioids depend on the test used and on the timing of the test relative to drug exposure, and they argue for the use of more than one test and both sexes in studies of this kind.

### Fentanyl and locomotor activity

Chronic fentanyl increased locomotor activity. In the SOF, the effect was significant during the first 5 min of the session, when exploration of the arena is highest, and only a trend over the full 15-min session, whereas in the LOF fentanyl increased the total distance traveled in both sexes. The groups did not differ in activity before pump implantation, so the post-implantation differences reflect the drug rather than pre-existing differences between the groups. Opioid-induced hyperactivity is a well-established phenomenon in rats that emerges as tolerance develops to the initial sedative effects and is mediated by increased mesolimbic dopamine transmission (Babbini and Davis, 1972; Di Chiara and Imperato, 1988). Consistent with this mechanism, fentanyl and related compounds stimulate locomotor activity and produce locomotor sensitization in rodents (Varshneya et al., 2019; Gaulden et al., 2021). The present findings show that this hyperactivity persists during continuous fentanyl exposure at a dose that produces dependence. Notably, fentanyl did not increase the distance traveled in the EPM. This is presumably because movement on the elevated maze is constrained by the aversiveness of the open arms, meaning the EPM was less affected by the locomotor-stimulant effect of fentanyl than the open field tests.

### Physical dependence and body weight

Naloxone precipitated a robust somatic withdrawal syndrome in fentanyl-treated rats, with a more than twentyfold increase in the total number of somatic signs, whereas naloxone had no effect in saline-treated rats. This confirms that the fentanyl regimen produced physical dependence and is consistent with previous work from our laboratory showing that continuous fentanyl administration produces dose-dependent affective and somatic withdrawal signs (Bruijnzeel et al., 2006; Bruijnzeel et al., 2007; Liu et al., 2008). Males and females did not differ in the total number of somatic signs or in any individual sign. This contrasts with the greater number of somatic signs that we observed in male than in female rats during precipitated nicotine withdrawal (Tan et al., 2019) and with the sex differences reported for fentanyl self-administration and relapse vulnerability (Townsend et al., 2019; Bakhti-Suroosh et al., 2021), and indicates that, at least for the somatic component of withdrawal from continuously delivered fentanyl, the two sexes are similarly affected. The presence of dependence in the same animals that displayed reduced anxiety-like behavior during active fentanyl exposure illustrates the opposing affective states associated with the presence and absence of the drug and is consistent with negative-reinforcement models of opioid use, in which drug taking is maintained both by the relief of aversive states and by the avoidance of withdrawal (Koob, 2020).

Chronic fentanyl also attenuated body weight gain, and this effect was larger in males than in females. Saline-treated males gained approximately 8% of their body weight over 13 days, whereas fentanyl-treated males did not gain weight and remained slightly below their initial post-operative weight. Reduced weight gain and weight loss are well-established consequences of chronic opioid administration (Gellert and Holtzman, 1978; Bruijnzeel et al., 2006; Bruijnzeel et al., 2007) and reflect, among other factors, opioid effects on food intake, gastrointestinal motility, and energy expenditure. The mechanisms underlying the greater sensitivity of males are not known but may include the larger growth trajectory of young adult males, which provides more room for a reduction in weight gain, and sex differences in the endocrine responses to opioids, including the suppression of the hypothalamic-pituitary-gonadal axis and gonadal steroids by chronic opioid exposure (Katz and Mazer, 2009; Vuong et al., 2010).

### Limitations and conclusions

Several limitations should be considered. First, behavioral testing was conducted while the osmotic minipumps continued to deliver fentanyl, so the anxiolytic-like effect was observed in the dependent state with the drug on board; anxiety-like behavior during spontaneous withdrawal, when the opposite effect would be expected (Schulteis et al., 1998; Fujii et al., 2019), was not assessed. Second, a single fentanyl dose was studied, and the dose-dependence of the anxiolytic-like and locomotor effects, as well as their persistence after fentanyl is discontinued, remains to be determined; chronic morphine, for example, impairs the extinction of conditioned fear a week after the last injection (Gu et al., 2008), so that lasting effects on anxiety-related behavior after the end of fentanyl exposure cannot be excluded. Third, the EPM and LOF tests were conducted at different times after pump implantation. Fourth, the sample size of eight rats per group was adequate to detect the main effects reported here but may have limited the power to detect smaller effects, particularly in females, and the estrous cycle was not monitored. Finally, because activity and anxiety-like behavior are related in exploration-based tests, an influence of locomotor activity on the anxiety-related measures, particularly in the LOF, cannot be excluded.

In conclusion, chronic fentanyl at a dose that produces physical dependence decreased anxiety-like behavior in both the EPM and the LOF, increased locomotor activity, and attenuated body weight gain in a sex-dependent manner. The anxiolytic-like effect was most pronounced in males in the EPM and in females in the LOF, which indicates that sex differences in the affective effects of opioids depend on the behavioral test and the stage of drug exposure. These findings add to the evidence that opioids reduce anxiety-like behavior during continuous administration (Fujii et al., 2019; Bruijnzeel et al., 2022), support the view that relief of anxiety contributes to the maintenance of opioid use (Martins et al., 2009; Koob, 2020), and underscore the importance of evaluating both sexes, using more than one test, and dissociating anxiety-like behavior from locomotor activity when characterizing the affective consequences of opioid exposure. These findings have significant clinical implications. Patients with chronic pain often suffer from comorbid anxiety (McWilliams et al., 2003; Asmundson and Katz, 2009), and the present findings suggest that chronic fentanyl administration may continue to provide relief from this anxiety despite the development of physical dependence and tolerance.

## Declarations

### Author contributions

L. Caglayan: Investigation, Formal analysis, Writing – Review & Editing. F. Zeyl: Investigation, Writing – Review & Editing. E. Gamba: Investigation, Writing – Review & Editing. M. Febo: Supervision, Writing – Review & Editing. A. Bruijnzeel: Conceptualization, Investigation, Formal analysis, Writing – Original Draft, Visualization, Supervision, Funding acquisition.

### Ethics approval

All procedures were approved by the University of Florida Institutional Animal Care and Use Committee and were conducted in accordance with the National Institutes of Health guidelines for the care and use of laboratory animals.

### Data availability

The datasets generated and analyzed during the current study, and the Python code used to generate the occupancy heatmaps, are available from the corresponding author on reasonable request.

### Declaration of generative AI use

During the preparation of this work the authors used Claude (Anthropic, San Francisco, CA, USA) to check grammar and improve the flow of the text, to write the SPSS syntax and the Python code with which the occupancy heatmaps (Figure 6) and the correlation matrices (Figure 8) were computed and plotted from the raw data, and to check the reference list. The authors reviewed and edited all output and take full responsibility for the content of the publication.

