## Supplemental tables for "Chronic fentanyl induces sex-specific anxiolytic-like effects in opioid-dependent rats"

**Supplement**

**Table S1.** Total distance traveled (cm) during the three pre-pump baseline sessions (days 1–3) in the small open field, by sex and drug treatment.

| **Group** | **Day 1** | **Day 2** | **Day 3** |
| --- | --- | --- | --- |
| ***Full 15-min session*** | | | |
| Male saline | 4,313 ± 431 | 3,897 ± 267 | 4,210 ± 344 |
| Male fentanyl | 4,583 ± 268 | 4,108 ± 287 | 4,441 ± 347 |
| Female saline | 5,480 ± 247 | 5,000 ± 373 | 4,958 ± 461 |
| Female fentanyl | 5,391 ± 201 | 4,736 ± 277 | 3,923 ± 245 |
| ***First 5-min block*** | | | |
| Male saline | 2,104 ± 103 | 1,988 ± 92 | 2,242 ± 156 |
| Male fentanyl | 2,140 ± 89 | 1,918 ± 69 | 2,273 ± 114 |
| Female saline | 2,351 ± 141 | 2,325 ± 147 | 2,420 ± 178 |
| Female fentanyl | 2,295 ± 127 | 2,171 ± 158 | 1,982 ± 117 |

Group means ± SEM (n = 8 per group).

**Table S2.** Total distance traveled (cm) in the small open field before and after pump implantation, by sex and drug treatment.

| **Group** | **Baseline** | **Post-pump** | **Post-pump, adjusted** |
| --- | --- | --- | --- |
| ***Full 15-min session*** | | | |
| Male saline | 4,140 ± 284 | 3,420 ± 381 | 3,716 ± 417 |
| Male fentanyl | 4,377 ± 272 | 3,951 ± 443 | 4,090 ± 402 |
| Female saline | 5,146 ± 301 | 3,852 ± 328 | 3,481 ± 428 |
| Female fentanyl | 4,683 ± 185 | 4,702 ± 531 | 4,638 ± 398 |
| ***First 5-min block*** | | | |
| Male saline | 2,111 ± 96 | 1,885 ± 170 | 1,913 ± 178 |
| Male fentanyl | 2,110 ± 74 | 2,239 ± 209 | 2,267 ± 178 |
| Female saline | 2,365 ± 88 | 1,913 ± 143 | 1,842 ± 187 |
| Female fentanyl | 2,150 ± 113 | 2,512 ± 180 | 2,525 ± 177 |

Observed group means ± SEM (n = 8 per group). Baseline is the mean of the three pre-pump sessions (days 1–3) and post-pump is the single session after pump implantation; these observed post-pump values are the ones plotted in Figure 2. The right-hand column gives the post-pump means after adjustment for baseline (estimated marginal means ± SE at the grand-mean covariate value) from the two-way (sex × drug) ANCOVA reported in the Results.
